# A ventral hippocampus to nucleus accumbens pathway regulates impulsivity

**DOI:** 10.1101/2025.11.18.689154

**Authors:** Molly E. Klug, Léa Décarie-Spain, Logan Tierno Lauer, Alicia E. Kao, Olivia P. Moody, Nicolas R. Morano, Haoyang Huang, Don B. Arnold, Emily E. Noble, Scott E. Kanoski

**Affiliations:** Human and Evolutionary Biology Section, Department of Biological Sciences, Dornsife College of Letters, Arts and Sciences, University of Southern California, USA; Molecular and Computational Biology Section, Department of Biological Sciences, Dornsife College of Letters, Arts and Sciences, University of Southern California, USA; Nutritional Sciences, College of Family and Consumer Sciences, University of Georgia, USA

**Keywords:** obesity, reward, addiction, impulse, craving, mesolimbic

## Abstract

Heightened impulsivity is attributed with substance abuse disorders, gambling, and obesity. The ventral hippocampus (vHPC) has recently been linked with impulse control, yet the neurobiological and behavioral mechanisms through which this control occurs are unknown. A subset of vHPC neurons project to the nucleus accumbens (ACB), a brain region known for regulating reward and motivation. Here, we evaluated the role of vHPC-ACB signaling in food-directed impulsivity using fiber photometry and transsynaptic chemogenetic and behavioral approaches. Male and female rats were trained in the differential reinforcement of low rates of responding (DRL) test of impulsive action, where they learn to withhold lever presses for 20 seconds to obtain a palatable food reward. Photometry recordings during DRL revealed analogous elevations in calcium-dependent activity in the vHPC and ACB during the 5s immediately prior to a non-impulsive vs. an impulsive response. Chemogenetic silencing of ACB-projecting vHPC neurons elevated impulsive responses in DRL relative to vehicle treatment in males but not females, yet had no effect on home cage food intake, operant-based motivation to work for palatable food, impulsive choice, or anxiety-like behavior. To determine whether this impulse control circuit requires vHPC->ACB communication independent of collateral targets of ACB-projecting vHPC neurons, we utilized a novel transsynaptic viral approach to selectively silence ACB neurons that receive synaptic glutamatergic signaling from the vHPC. Results reveal that inhibition of ACB neurons receiving vHPC signaling elevates impulsive action in the DRL task relative to vehicle treatment. Collective results reveal a hippocampal-striatal circuit that regulates impulsive action in males.

## INTRODUCTION

Impulsivity, characterized by acting with little-to-no forethought for action consequences, has been implicated in obesity, binge-eating disorder (BED), compulsive gambling, and various substance abuse disorders ^1–9^. One hypothesis for the etiology of the obesity crisis, which affects over 40% of adults in the US^10^, centers around deficits in behavioral inhibition and increased reward-salience, which in an obesogenic environment can lead to impulsive food-directed behaviors resulting in excessive caloric consumption ^5,11,12^. Consistent with this hypothesis, obesity is associated with lower inhibitory control scores, an outcome predictive of success in behavioral weight loss intervention programs ^1^. Further, deficits in behavioral inhibition and impulse control are strongly linked with BED, as well as both childhood and adult obesity ^1,2,13,14^.

Impulsivity, while a complex and multi-faceted personality trait, can be divided into two distinct subtypes: behaviors resulting from a failure to suppress an inappropriate action (impulsive action) and behaviors wherein the choice between two or more options is made without proper consideration of future consequences (impulsive choice). While the neural circuitry regulating impulse control is poorly understood, emerging evidence indicates that impulsive actions and impulsive choices are modulated by both overlapping and distinct neural substrates ^15,16^. Furthermore, there is significant support for sexual dimorphism in trait impulsivity, with males being generally more susceptible than females ^17–20^. Thus, understanding the neural substrates underlying impulsivity must involve both behavioral task- and sex-specific evaluations.

The hippocampus (HPC), a brain region crucial for the encoding, consolidation, and retrieval of episodic and spatial memories, has recently been linked in regulating impulse control of behaviors directed towards obtaining palatable foods ^21,22^. The HPC is also implicated in the control of eating behavior more generally, as HPC damage is associated with impaired satiety^23,24^ and selective HPC lesions in rodents leads to increased food intake and body weight gain ^25^. Further, acute inactivation of either the dorsal or ventral HPC subregion immediately after a meal increases subsequent food consumption in rodents ^26,27^, and an orexigenic hippocampal subnetwork associated with dysregulated eating and body mass index in humans was recently identified ^28^. However, the neural systems through which the hippocampus regulates impulsive responding for palatable foods remain poorly understood.

A putative downstream site-of-action for HPC-mediated impulse control is the nucleus accumbens (ACB) ^21^. The ACB is associated with reward processing and controlling impulsivity^29–32^, and ACB-projecting hippocampal neurons enhance food palatability in rodents ^33^. Further, a recent pilot study in humans revealed that deep brain stimulation in the ACB for patients with severe BED and obesity improved loss-of-control eating frequency and was associated with weight-loss ^34^. While previous research on HPC-to-ACB circuitry has focused its role in neuropsychiatric disorders ^35,36^, contextual conditioning ^37,38^, motivational conflict ^39^, and on non-food related reward processing (e.g., drugs of abuse) ^40,41^, we hypothesized that HPC neuronal projections to the ACB, more specifically, ventral HPC (vHPC; field CA1) inputs to the ACB shell subregion (ACBsh), regulate impulsive responding for palatable foods. This hypothesis is evaluated in present experiments in both male and female rats, and using tests of both impulsive action and impulsive choice as well as complementary transsynaptic functional neural circuit disconnection approaches.

## METHODS and MATERIALS

### Animals

Male and female Sprague-Dawley rats (Envigo, Indianapolis, IN; postnatal day [PND] 60-70; 250-275g on arrival for males, 180-200g on arrival for females) were individually housed in a temperature-controlled vivarium with *ad libitum* access (except where noted) to water and food (LabDiet 5001, LabDiet, St. Louis, MO) on a 12h:12h reverse light/dark cycle. All procedures were approved by the Institute of Animal Care and Use Committee at the University of Southern California.

### Estrus tracking

To identify estrous stage, vaginal cytology was investigated by vaginal lavage ^42–45^. Cytology samples were collected in 0.9% sterile saline (Teknova) 1.5 hr prior to the start of the dark cycle. Vaginal smears were examined at 10x using a Nikon 80i (Nikon DS-QI1,1280X1024 resolution, 1.45 megapixel). Stages were assigned at the same time each day to minimize the incidence of transitional and ‘missed’ stages and were based on vaginal cell morphology (presence or absence of epithelial cells, cornified cells, and leucocytes) as described in ^42^.

### Surgical procedures

For all surgical procedures, rats were anesthetized and sedated via intramuscular injections of ketamine (90 mg/kg), xylazine (2.8 mg/kg), and acepromazine (0.72 mg/kg). Rats were also given analgesic (subcutaneous injection of ketoprofen [5mg/kg]) after surgery and once daily for 3 subsequent days thereafter. All rats recovered for at least one-week post-surgery prior to experiments.

### Viral injections

For stereotaxic injections of viruses and tracers, rats were first anesthetized and sedated with a ketamine (90 mg/kg)/xylazine (2.8 mg/kg)/acepromazine (0.72 mg/kg) cocktail. Animals were shaved, surgical site was prepped with iodine and ethanol swabs, and animals were placed in a stereotaxic apparatus for stereotaxic injections. Viruses were delivered using a microinfusion pump (Chemyx, Stafford, TX, USA) connected to a 33-gauge microsyringe injector attached to a PE20 catheter and Hamilton syringe. Flow rate was calibrated and set to 5 µl/min; injection volume was 200 nl/site. Injectors were left in place for 2 min postinjection. Following injections, animals were either sutured or surgically implanted with a cannula. All experimental procedures occurred 21 days post virus injection to allow for transduction and expression. Successful virally mediated transduction was confirmed postmortem in all animals via IHC staining using immunofluorescence-based antibody amplification to enhance the fluorescence followed by manual quantification under epifluorescence illumination using a Nikon 80i (Nikon DS-QI1,1280X1024 resolution, 1.45 megapixel).

For recording of CA2+ activity, 400nl of AAV9-hSyn-GCaMP7s-WPRE (Cat. no. 104487-AAV9, Addgene) was unilaterally injected at the following coordinates: CA1v: −4.9 mm anterior/posterior (AP), +-4.8 mm medial/lateral (ML), −7.8 mm dorsal/ventral (DV) (0 reference point at bregma for ML, AP, 0 reference point at skull site for DV) ACB: +1.2 mm anterior/posterior (AP), +-1.0 mm medial/lateral (ML), −6.75 mm dorsal/ventral (DV) (0 reference point at bregma for ML, AP, 0 reference point at dura for DV). An optic fiber was implanted at the injection site as described below.

For chemogenetic inhibition of CA1v neurons projecting to the ACB via DREADDS, an AAV2-hSyn-DIO-hM4D(Gi)-mCherry, (DIO-DREADDs; Cat. no. 44362-AAV2, Addgene) was bilaterally injected at the following coordinates: −4.9 mm anterior/posterior (AP), +-4.8 mm medial/lateral (ML), −7.8 mm dorsal/ventral (DV) (0 reference point at bregma for ML, AP, 0 reference point at skull site for DV). Additionally, an AAV2(retro)-hSYN1-eGFP-2A-iCre-WPRE; (Retro-cre; Cat. no. VB4855, Vector Biolabs) was bilaterally injected at the following coordinates, to allow for targeted inhibition of ACB-projecting CA1v neurons: +1.2 mm anterior/posterior (AP), +-1.0 mm medial/lateral (ML), −6.75 mm dorsal/ventral (DV) (0 reference point at bregma for ML, AP, 0 reference point at dura for DV). The injection volume was 300 nl/site.

For pathway-specific chemogenetic inhibition of ACB neurons receiving glutamatergic projections from the CA1v, AAV8-ATLAS_cre_ and pAAV-DIO0BACE-HA (ATLAScre; Gift from the Arnold Lab ^46^; available on Addgene, ID 232351 and ID232353) was bilaterally injected at a 1:1 ration in the following coordinates: −4.9 mm anterior/posterior (AP), +-4.8 mm medial/lateral (ML), −7.8 mm dorsal/ventral (DV) (0 reference point at bregma for ML, AP, 0 reference point at skull site for DV). Additionally, AAV2-hSyn-DIO-hM4D(Gi)-mCherry, (DIO-DREADDs; Cat. no. 44362-AAV2, Addgene) was bilaterally injected at the following coordinates: +1.2 mm anterior/posterior (AP), +-1.0 mm medial/lateral (ML), −6.75 mm dorsal/ventral (DV) (0 reference point at bregma for ML, AP, 0 reference point at dura for DV). The injection volume was 300-400 nl/site.

### Fiber photometry

Surgeries were adapted from our procedures described previously ^47–50^. Rats were first anesthetized and sedated via intramuscular injection of a ketamine (90 mg/kg), xylazine (2.8 mg/kg), and acepromazine (0.72 mg/kg) cocktail, prepped for surgery, and placed in stereotaxic apparatus. They were given subcutaneous injections of buprenorphine SR (0.65 mg/kg) during surgery as an analgesic. A fiber-optic cannula (Doric Lenses Inc, Quebec, Canada; flat 400-μm core, 0.48 numerical aperture [NA]) was implanted in the CA1v and ACB at the following coordinates: CA1v: −4.9 mm anterior/posterior (AP), +-4.8 mm medial/lateral (ML), −7.8 mm dorsal/ventral (DV) (0 reference point at bregma for ML, AP, 0 reference point at skull site for DV) ACB: +1.2 mm anterior/posterior (AP), +-1.0 mm medial/lateral (ML), −6.75 mm dorsal/ventral (DV) (0 reference point at bregma for ML, AP, 0 reference point at dura for DV). The optic fibers were then affixed to the skull with jeweler’s screws, instant adhesive glue, and dental cement. All subjects were given 1 week to recover from surgery prior to experiments.

In vivo fiber photometry was performed according to previous work ^47–50^. Photometry signal was acquired using the Neurophotometrics fiber photometry system (Neurophotometrics, San Diego, CA) at a sampling frequency of 40 Hz and administering alternating wavelengths of 470 nm (Ca^2+^ dependent) or 415 nm (Ca^2+^ independent). The fluorescence light is transmitted through an optical patch cord (Doric Lenses) and converges onto the implanted optic fiber, which in turn sends back neural fluorescence through the same optic fiber/patch cord and is focused onto a photoreceiver. All behavioral responses (i.e. lever presses) were time-stamped using the data acquisition software (Bonsai). The resulting signals were then corrected by subtracting the Ca^2+^ independent signal from the Ca^2+^ dependent signal to calculate fluorescence fluctuations due to Ca^2+^ (corrected signal) and not due to baseline neural activity or motion artifacts and fitted to a biexponential curve. The corrected fluorescence signal was then normalized within each rat by calculating the ΔF/F using the average fluorescence signal for the entire recording and converting the signal to *z*-scores. The normalized signal was then aligned to behavioral events and data extraction was done using original MatLab code.

### Drug preparation

For chemogenetic inhibition, 1 ml/kg DCZ (100 µg/kg) or vehicle (1% DMSO in 99% saline) is administered intraperitoneally through a syringe, 5 mins prior to behavioral testing. Doses and concentration of DCZ and vehicle were based on previous work ^48^.

### Differential reinforcement of low rates of responding

As previously described ^21,51^, minimally-food restricted rats (food removed from home cage 1hr before dark onset) were trained in the early nocturnal phase on an operant lever-pressing task to obtain a single, high-fat high-sugar 45mg pellet (HFHS; F05989, Bio-serv; 35% kcal fat and sucrose-enriched) for each press. Training and test sessions were conducted in operant conditioning boxes (Med Associates Inc, St. Albans, VT). After progressively increased delay periods (0-, 5-, 10-second delays, 5 days of training for each schedule) over four weeks of training, rats are given a ‘DRL-2o’ schedule for ten days in which they must wait 20 seconds between lever presses for reinforcement. A lever press prior to conclusion of the 20-second interval resets the clock, is not reinforced, and is considered an impulsive response. On the other hand, a lever press after the conclusion of the 20-second interval is reinforced, and is considered a nonimpulsive response. Testing took place on the DRL20 schedule at least 2 days after the completion of training. The periods of 5 seconds immediately before and after a lever press was determined a priori for analyses of calcium-dependent activity associated with a non-impulsive (reinforced) vs. an impulsive (non-reinforced) press.

### Delay discounting procedure

As previously described ^21^, rats were trained in a series of 4 blocks of 5 choice trials (delay training), wherein one lever consistently delivers one pellet and the other delivers 4 pellets in a time delay of 0s, 5s, 10s, and 20s corresponding to blocks 1, 2, 3, and 4. Each block began with a forced trial, in random order, during which a stimulus light above each lever was lit. During choice trials levers remained extended for 10s or until a lever response is made, after which the levers were retracted, and the stimulus light remained illuminated until the delivery of the pellet. Training and test sessions were conducted in operant conditioning boxes (Med Associates Inc, St. Albans, VT). Delay training lasted for a total of 24 days, after which testing took place at least 2 days after training.

### BioDAQ food monitoring system and meal pattern analysis parameters

Rats were individually housed in a BioDAQ Food and Water intake monitoring system (Research Diets Inc. New Brunswick, NJ) which recorded episodic ad libitum food intake activity from rats in their home cages. All peripheral sensor controllers (PSCs) were validated for accuracy using 10.00g standard weights with an allowable margin of error +/- 0.05g prior to the start of the experiment as well as before test days. All PSCs were required to have QUIET readings (i.e., no movement at food hopper) prior to the start of any session. BioDAQ data was analyzed using an inter meal interval (IMI) of 900 seconds to separate meals, and bouts were filtered to be between -9.00 and 9.00g in size. Data was also set to have a period that started at the onset of the dark cycle when the behavioral task would begin, in this case 11:00AM to coincide with the light schedule in the room in which they were housed. The number of periods in the day was set to 24, resulting in 1hr long period bins from which cumulative intake as well as average meal size and duration could be calculated. Following exportation to Microsoft Excel, values were obtained from the “PSC by period” tab to record individual animals’ intake.

### Progressive Ratio task (PR)

Rats were trained to lever press for 45mg pellet (HFHS; F05989, Bio-serv; 35% kcal fat and sucrose-enriched) in operant conditioning boxes over the course of 6 days, with a 1hr session each day. The first 2 days consisted of fixed ratio 1 with <u>autoshaping</u>, wherein animals would receive 1 pellet for each correct lever press (FR1), and a pellet would dispense automatically every 10 min. The next 4 days consisted of FR1 without autoshaping (2 days) and FR3 (2 days), wherein animals would receive 1 pellet for every 3 presses on the active lever. On the test day rats were placed back in the operant chambers to lever press for HFHS pellets under a progressive ratio reinforcement schedule. The response requirement increased progressively using the following formula: F(i) = 5ê0.2i-5, where F(i) is the number of lever presses required for the next pellet at i = pellet number and the breakpoint was defined as the final completed lever press requirement that preceded a 20-min period without earning a reinforcer, as described previously ^47,52^ .

### Zero Maze task (ZM)

Following established procedures ^49^, the zero maze apparatus was used to examine exploratory and anxiety-like behavior. The apparatus consisted of an elevated circular track (11.4cm wide track, 73.7cm height from track to ground, 92.7cm exterior diameter) that is divided into four equal length segments: two sections with 3-cm high curbs (open) and two sections with 17.5-cm height walls (closed). Ambient lighting was used during testing. Rats were placed in the maze on an open section of the track and allowed to roam freely for 5 min. The apparatus was cleaned with 10% ethanol between rats. The total distance travelled and time spent in the open segments of the apparatus were measured via video recording using ANY-maze activity tracking software.

### Immunohistochemistry

Rats were anesthetized and sedated with a ketamine (90 mg/kg)/xylazine (2.8 mg/kg)/acepromazine (0.72 mg/kg) cocktail, then transcardially perfused with 0.9% sterile saline (pH 7.4) followed by 4% paraformaldehyde (PFA) in 0.1 M borate buffer (pH 9.5; PFA). Brains were dissected out and post-fixed in PFA with 15% sucrose for 24 h, then flash frozen in isopentane cooled in dry ice. Brains were sectioned to 30-µm thickness on a freezing microtome. Sections were collected in 5 series and stored in antifreeze solution at −20 °C until further processing.

General fluorescence IHC labeling procedures were performed for histological confirmation. The following antibodies and dilutions were used: rabbit anti-RFP (1:2000, Rockland Inc., Limerick, PA, USA), and chicken anti-GFP (1:1000, Cat. no. ab13970, Abcam, Cambridge, UK), ALFA-TAG (1:1000, Cat No: N1502, NanoTag Biotechnologies, Gottingen, Germany). Antibodies were prepared in 0.02 M potassium phosphate-buffered saline (KPBS) solution containing 0.2% bovine serum albumin and 0.3% Triton X-100 at 4 °C overnight. After thorough washing with 0.02 M KPBS, sections were incubated in secondary antibody solution. All secondary antibodies were obtained from Jackson Immunoresearch and used at 1:500 dilution at 4 °C, with overnight incubations (Jackson Immunoresearch; West Grove, PA, USA). Sections were mounted and coverslipped using 50% glycerol in 0.02 M KPBS and the edges were sealed with clear nail polish. Photomicrographs were acquired using either a Nikon 80i (Nikon DS-QI1,1280X1024 resolution, 1.45 megapixel) under epifluorescence or darkfield illumination.

### Statistical Analyses

All statistical analyses were conducted using GraphPad Prism 10.0.3 (GraphPad Software, Inc. Boston, Massachusetts, USA). Data are reported as mean ± SEM. T-tests were used to analyze AUC deltas, DRL, 1- and 6hr BioDAQ measures in males and females respectively, PR, and ZM. Two-way ANOVA was used to analyze DD, and a mixed-effects analysis was used to analyze all BioDAQ data from 0-6hrs in males. A one-way ANOVA was used to analyzed BioDAQ data in females when broken down by estrus cycle. Simple linear regression was used to analyze the correlation between DRL efficiency and change in photometry signal. Statistical significance was set at a p-value ≤ 0.05.

## RESULTS

### Calcium-dependent activity in the vCA1 and ACB predicts control of impulsive action

To examine how activity in the vCA1 and ACB relates to impulsive responding for palatable foods, bulk calcium-dependent activity was recorded from both the CA1v and ACB during the DRL task, as described above (Figure 1A). Successful viral transfusion and cannula placement were confirmed via histology for both the CA1v and ACB (Figure 1B). Animals successfully learned the DRL task (Figure 1C), reaching ∼60% efficiency in the task (Figure 1D), consistent with our previous work ^21^. During the DRL test recording session, calcium-dependent activity in the ACB was significantly higher in the 5s preceding a reinforced (i.e. non-impulsive) response, relative to an impulsive (i.e. non-reinforced) response (Figure 1E, F). Calcium-dependent activity in the vCA1 was similarly elevated in the 5s preceding a reinforced response, relative to an impulsive response (Figure 1G, H). Additionally, calcium-dependent activity in both brain regions was lowered in the 5s following a reinforced response relative to an impulsive response (Supp. Figure 1A, B). Efficiency in the DRL task was significantly correlated with the difference in calcium-dependent activity between a non-impulsive and an impulsive response, both before and after a lever press (Supp. Figure 1C, D). These data support the feasibility that CA1v projections to the ACBsh are functionally involved in regulating impulsive actions directed towards palatable food reinforcement.

**Figure 1.**
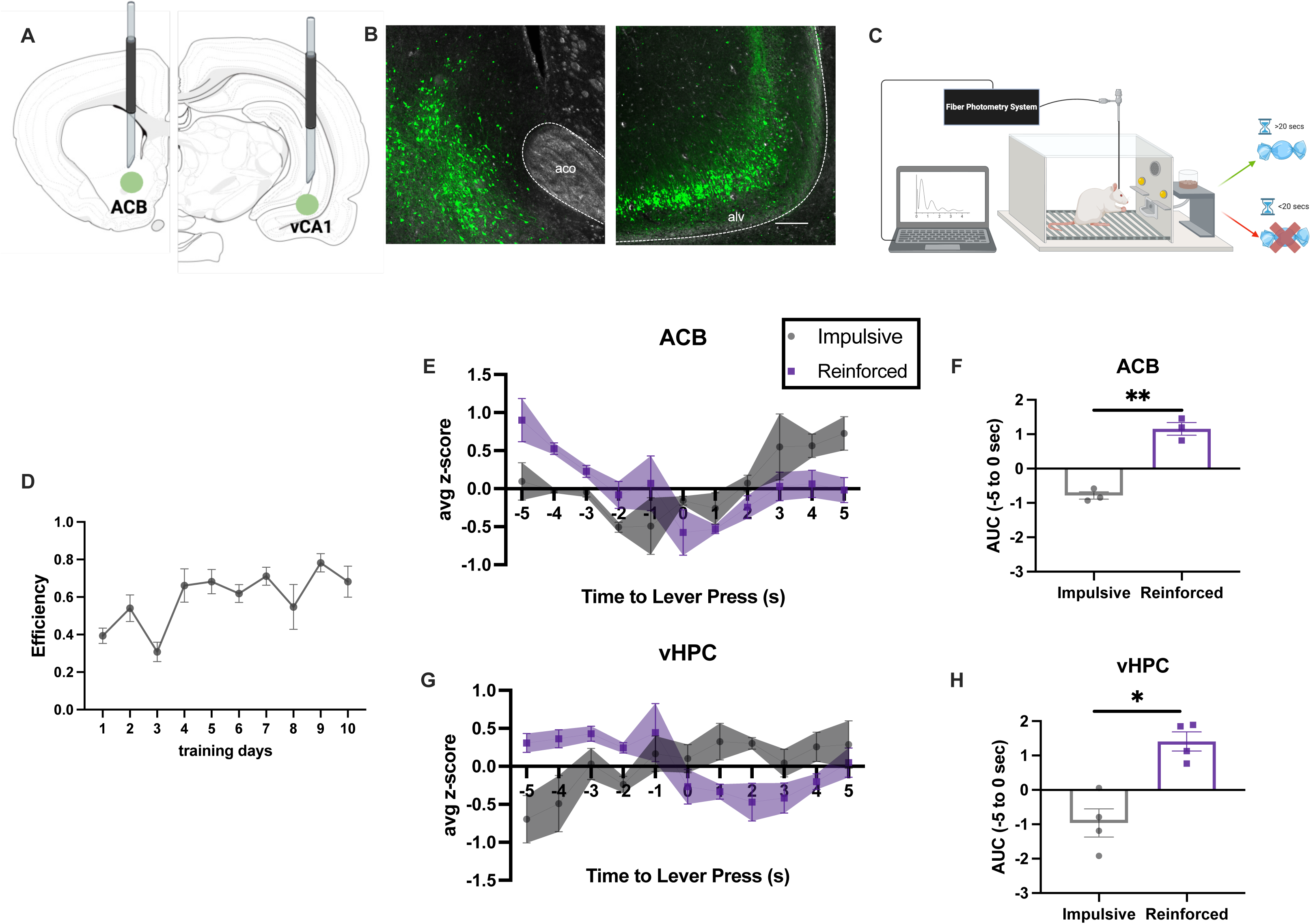
Calcium-dependent activity is increased in both the vHPC and ACBsh prior to a non-impulsive response in the DRL task. **(A)** Diagram depicting surgical procedure for fiber photometry recordings of Ca^2+^-dependent activity in vHPC and ACBsh.**(B)** Representative images of surgical targets for the ACBsh (left) and vHPC (right) (Scale bar, 200 um). **(C)** Diagram depicting the DRL task with photometry recording. Ca^2+^-dependent activity is elevated in the 5s before a non-impulsive lever press, relative to an impulsive press in the vHPC **(D-F)** and ACBsh **(G-I)**. (vHPC n = 4, ACBsh n = 3; all between-subjects design; Data are means ± SEM; *p<0.05, ***p<0.001).

### Silencing of ACB-projecting vCA1 neurons increases impulsive action in males, but not impulsive choice in either sex

To directly test the CA1v-ACB pathway’s involvement in impulsive responding, animals were injected with a retrograde cre-expressing virus in the ACB and a cre-dependent inhibitory DREADDs in the CA1v, an approach that selectively targets CA1v neurons that project to the ACB for reversible chemogenetic inhibition (Figure 1A). Successful viral transfection was confirmed via IHC as describe above (Figure 2B). Animals were trained in the DRL task of impulsive action (Figure 2C). Animals successfully learned the task, reaching between 50-60% efficiency (Figure 2D, H). On test days, animals received intraperitoneal (IP) injections of either Deschloroclozapine (DCZ; DREADDs ligand) or vehicle (1% DMSO in 99% saline) to selectively inhibit ACB-projecting CA1v neurons. Results revealed that males receiving DCZ made significantly more responses on the active lever relative to animals receiving vehicle, while the total number of rewards earned was unaffected (Figure 2E, F), resulting in decreased efficiency in males receiving DCZ relative to vehicle (i.e., increased impulsive action; Figure 2G). However, females showed no change in task efficiency, lever presses, or rewards earned following DCZ treatment relative to vehicle (Figure 2I-K), even when controlling for the effects of estrus cycle stage (Supp. Figure 2D-F).

**Figure 2.**
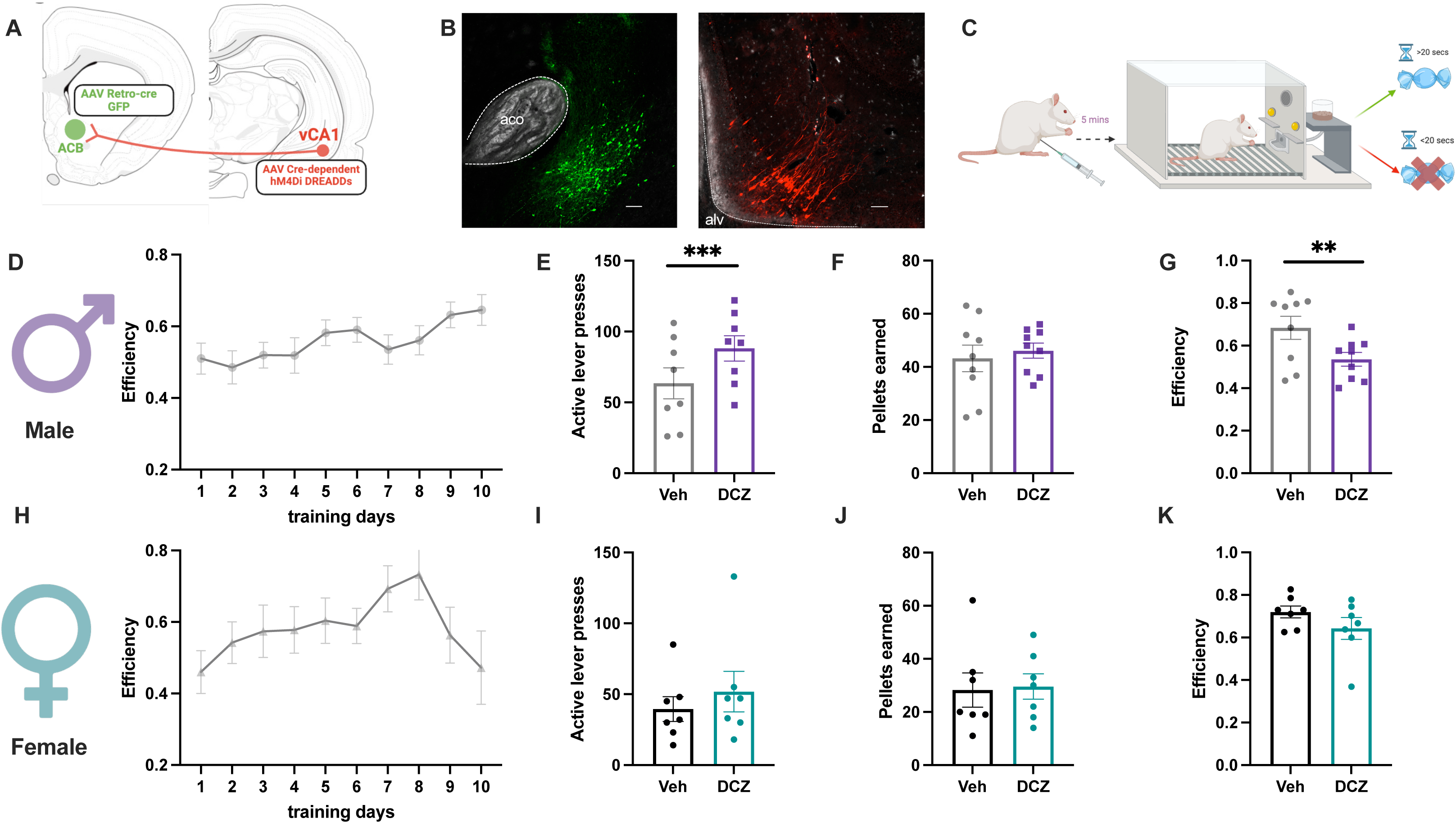
Silencing ACBsh-projecting CA1v neurons increases impulsive action in male rats. **(A)** Diagram depicting surgical procedure for chemogenetic inactivation of the vHPC-ACBsh pathway. **(B)** Representative images of surgical targets for the ACBsh (left) and vHPC (right) (Scale bar, 100um). **(C)** Diagram depicting the DRL task. DREADD inhibition of the vHPC-ACBsh pathway increases impulsive responding in the DRL task in males **(D-F)**, but not females **(G-I).**

As a control to ensure that DCZ administration itself was not playing a role in impulsive responding, a separate group of male rats was injected with cre-dependent inhibitory DREADDs in the CA1v, but were not given a retrograde, cre-expressing virus in the ACB, rendering the DREADDs nonfunctional. Administration of DCZ prior to the DRL task had no effect on efficiency, total lever presses, or rewards earned in this group (Supp. Figure 2I-K).

While the full neural circuitry regulating impulsive responding is poorly understood, evidence suggests that impulsive actions and impulsive choices are modulated by both overlapping and distinct neural substrates^15,21^. DRL is an operant test for impulsive actions, but fails to address questions of impulsive choices. Therefore, to test whether inhibition of the CA1v-to-ACB pathway differentially affects impulsive actions and impulsive choices, we used the same viral approach described above, paired with a delay discounting task (Supp. Figure 2A). Delay discounting refers to tendency to devalue a reward as the time delay associated with delivery of the reward increases. During testing, animals of both sexes exhibited standard discounting behavior, exhibited as increased responding on the immediate lever as the time delay on the larger reward lever progressively increased. However, no overall differences were found between DCZ- and vehicle-treated animals in males (Supp. Figure 2B) or females (Supp. Figure 2C, G), indicating that the CA1v-to-ACB pathway does not regulate impulsive choice (Supp. Figure 2B, C). Taken together, these results show that reversible inhibition of ACB-projecting CA1v neurons affects impulsivity in a sex- and task-dependent manner, thus further emphasizing the importance of evaluating sex as a variable for research on impulsivity, as well as the fact that impulsive action and choice are mediated by distinct neural circuitry.

### Silencing of ACB-projecting vCA1 neurons does not affect food intake, motivation to work for palatable food, or anxiety-like behavior

To test whether our results might be influenced by behavioral changes other than impulsivity, we performed a variety of behavioral assays to determine the effects of reversible inhibition of the CA1v-ACB pathway on food intake, reward motivation, and anxiety-like behavior. Given that our phenotype for impulsivity was not evident in females, these experiments were performed predominantly in male rats. Inhibition of the CA1v-ACB pathway did not affect cumulative intake of standard chow (Figure 3A; Supp. Figure 3A, G, J), nor did it affect the number of meals eaten (Figure 3B; Supp. Figure 3B, H, K) or the average meal size (Figure 3C; Supp. Figure 3C, I, L) for either males or females over a 1hr, 2hr, 3hr, 5hr, or 6hr period. Inhibition of this pathway also failed to affect these same parameters when animals were given a high-fat, high-sugar (HFHS) chow (Figure 3D-F; Supp Figure 3D-F). Overall reward motivation was not affected by the CA1v-ACB pathway inhibition in the progressive ratio task in male rats (Figure 3H-J), nor was anxiety-like or exploratory behavior in the zero maze test (Figure 3K-M).

**Figure 3.**
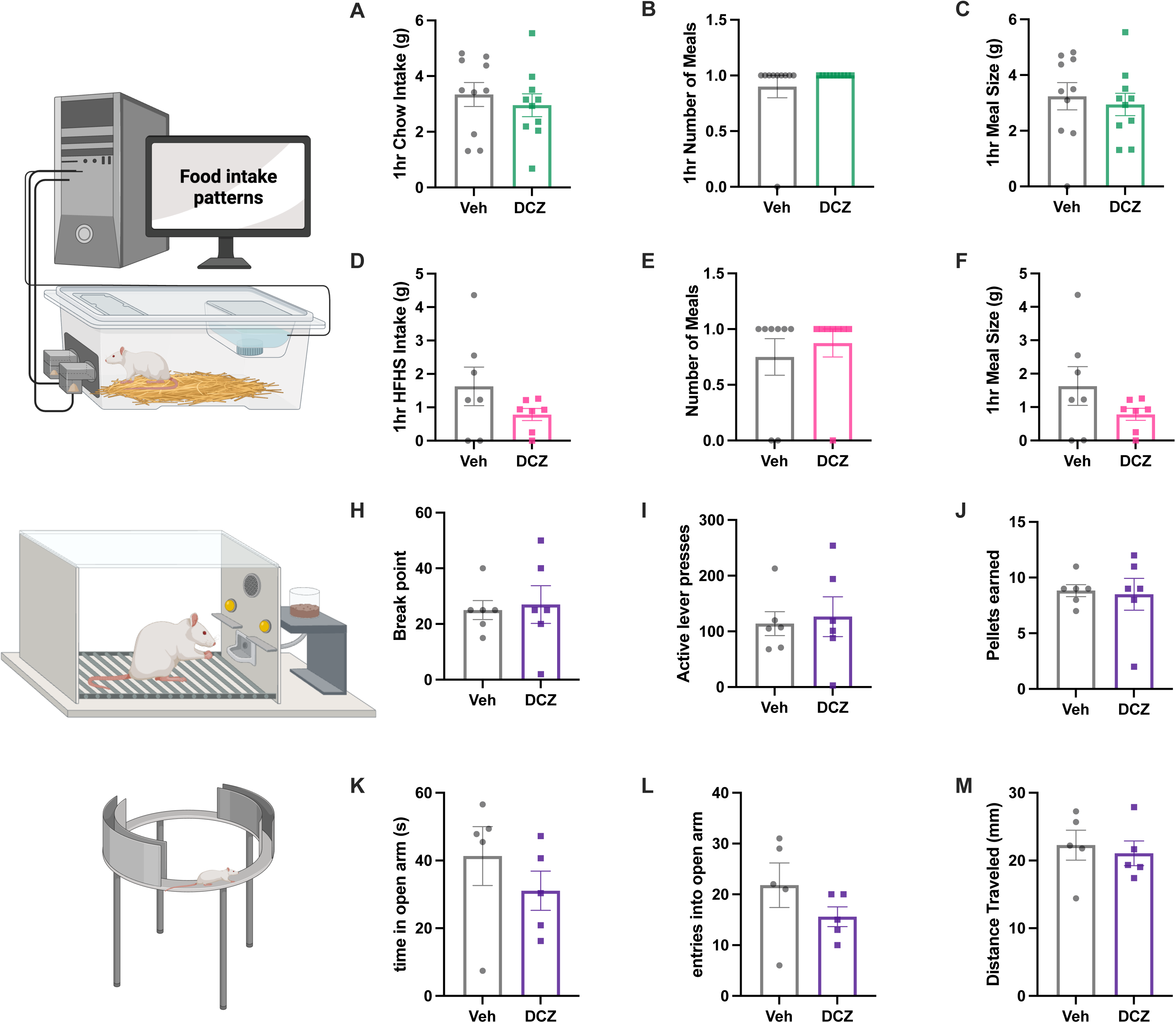
Silencing ACBsh-projecting CA1v neurons does not affect home cage intake of standard chow, HFHS chow, reward motivation, or anxiety-like behavior. Home cage intake and meal pattern parameters were unaffected by chemogenetic silencing for both standard chow (**A-C**) and HFHS chow (**D-F**). Reward motivation, as measured in the PR task, was unaffected by pathway inhibition (**H-J**), as was anxiety like behavior (**K-M**). (Standard chow n = 10, HFHS chow n = 8, PR n = 7, zero maze n = 5; within=subjects for food intake and reward motivation, between-subjects for anxiety-like behavior; Data are means ± SEM; *p<0.05, ***p<0.001).

### Increased impulsive action in males is specific to CA1v-ACB synaptic communication

Given that ventral HPC CA1 neurons are known to have collateral projections to various hypothalamic, striatial, septal, and cortical targets ^47,50^, we tested whether our results might be driven by collateral projections from ACB-projecting CA1v neurons, vs. direct CA1v->ACBsh signaling. We utilized a novel viral approach developed to target downstream neurons in a strictly anterograde, monosynaptic, and glutamatergic synaptic activity-dependent manner (ATLAS) ^46^. We paired this novel transsynaptic viral approach in the CA1v – in this case to drive cre recombinase expression downstream of CA1v glutamatergic terminals – with a cre-dependent inhibitory DREADDs in the ACB, to effectively isolate the CA1v-ACB-specific pathway for reversible silencing (Figure 4A, B). Results reveal that animals receiving DCZ made more presses on the active lever (Figure 4E), while earning a similar number of rewards (Figure 4F) compared to animals receiving vehicle. While this difference was not statistically significant, it nevertheless contributed to a significant decrease in efficiency in the DRL task – the primary measure of impulsive action in this test – when animals received DCZ relative to when they received the vehicle (Figure 4G). These results indicate that our earlier findings revealing that chemogenetic inhibition of ACB-projecting CA1v neurons increases impulsivity in this task were not due to inhibition of collateral projections of these neurons, but rather, were driven by inhibition of Ca1v to ACB glutamatergic signaling.

**Figure 4.**
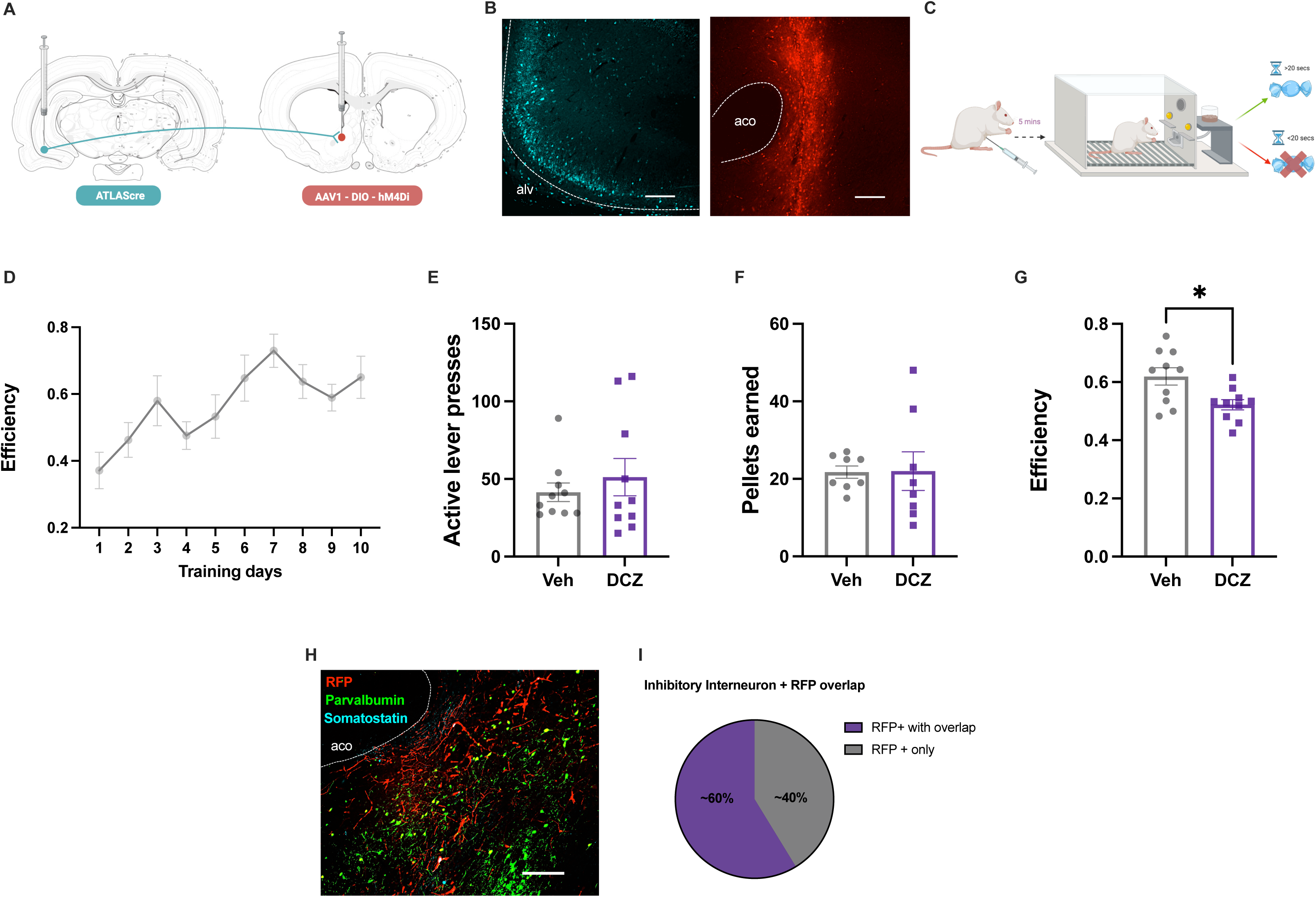
Increased impulsive action in males is specific to CA1v to ACB signaling. **(A)** Diagram depicting surgical procedure for chemogenetic inhibition of ACB neurons which receive projections from the CA1v. **(B)** Representative images of surgical targets for the ACBsh (left) and vHPC (right) (Scale bar, 100um). **(C)** Diagram depicting the DRL task. **(D)** Training efficiency for the DRL task. **(E-G)** Chemogenetic inhibition of ACB-projecting CA1v neurons increases impulsive responding in males. **(H, I)** A majority of ACB neurons receiving projections from the CA1v are inhibitory interneurons (For DRL: n = 10, within-subjects; For IHC n=3; Data are means ± SEM; *p<0.05).

### CA1v neurons target inhibitory interneurons in the ACB

In order to gain insight into the phenotype of ACB neurons receiving projections from the CA1v, we preformed IHC to co-stain for ACB neurons that receive synaptic input from CA1v neurons and the inhibitory interneuron markers, parvalbumin and somatostatin. Results show that roughly 60% of the neurons in the ACB which receive projections from the CA1v are inhibitory interneurons (Figure 4H, I), expressing either somatostatin or parvalbumin. These results indicate that HPCv->ACBsh regulation of impulsivity in males may predominantly involve modulation of intrinsic ACBsh activity via interneuron inhibitory GABAergic signaling.

## DISCUSSION

Palatable food-directed impulsivity is strongly implicated in obesity and BED ^1,2,5,6^, and this association may be driven by neural circuits that connect the hippocampus (HPC) with the nucleus accumbens (ACB) ^34,53,54^. Present results reveal that calcium-dependent activity in both the ACB (shell region) and the ventral HPC (vHPC; CA1v region) is significantly elevated prior to a non-impulsive lever press relative to an impulsive press in the operant DRL test of impulsive action, suggesting that each region may play a functional role in regulating impulse control. Consistent with these findings, reversible chemogenetic silencing of CA1v that project to the ACB resulted in significantly decreased efficiency in the DRL task, an outcome indicative of increased impulsive action. However, this result was only seen in males as chemogenetic inhibition of the CA1v-ACB pathway in females did not impact DRL performance, regardless of estrus stage. Furthermore, we found that chemogenetic inhibition of the CA1v-ACB pathway did not affect impulsive choice, as measured using the delay discounting task, in either male or females, nor did it affect home cage consumption of standard or HFD chow, motivation to work for palatable food, or anxiety-like behavior. By combining a novel transsynaptic glutamatergic synapse-based ATLAS approach ^46^ with chemogenetic silencing and immunostaining, we reveal that the CA1v-ACB pathway’s control of impulsive behavior are indeed pathway specific to CA1v synapses on ACB neurons and not collateral targets of these neurons. Additionally, we find that CA1v neurons communicate predominantly to inhibitory interneurons within the ACB.

Both the vHPC and ACB have recently emerged as likely substrates in regulating impulsive responding ^21,22,29–32^, and thus we sought to investigate the role of the CA1v-ACB pathway in mediating multiple forms of impulse control, looking at both impulsive actions and impulsive choices. Previous research suggests impulsive action and impulsive choice are dissociable behaviors ^15,55–57^, with studies finding that outcomes in impulsive choice and action measures are not correlated in either humans or rats ^16^. The vHPC has previously been implicated in the regulation of both impulsive actions ^21,52,58^ and impulsive choices ^59–62^, with vHPC loss-of-function approaches leading to increases in impulsive responding. While our results revealing that vHPC-ACB pathway inhibition increases impulsive actions are consistent with previous literature, our lack of effect on impulsive choice warrants follow-up consideration. Given the pathway-specific nature of the current study by using two distinct but complementary disconnection approaches, it is difficult to draw direct comparisons to previous research that targeted this pathway with less specificity. For example, Abela et al. investigated the role of the vHPC-ACB pathway in regulating impulsive choices using an excitotoxic contra-lesional approach, finding a significant difference in delay discounting behavior in animals with a disconnected vHPC-ACB pathway ^60^. However, contra-lesional approaches are not pathway-specific, and disrupt collateral projection targets to and from both regions. Further, these approaches are chronic and not reversible, and are thus vulnerable to compensatory neural adaptations. Therefore, our lack of an observed effect on impulsive choice behavior is likely based on the increased sensitivity of our pathway-specific manipulations compared to previous lesion-based literature. Taken together with previous literature, our results indicate that the influence of vHPC signaling on impulsive choice may be based on projections to targets other than the ACB (e.g., lateral hypothalamus or medial prefrontal cortex).

While we found that inhibition of ACB-projecting CA1v neurons increased impulsive action in males, we found no effect on either impulsive action or impulsive choice in females. Research investigating sex differences in impulsive responding is mixed, with outcomes in humans varying depending on the population and task being studied ^19,20,63^. In animal models, outcomes can vary based on the developmental timing of testing ^17,18,20,64^, though males tend to exhibit higher impulsivity in impulsive actions ^18,20,65,66^ whereas females are more prone to impulsive choices ^20^. Our current results are largely consistent with these findings. However, while previous studies have shown a role of estrus/menstrual cycle in the regulation of impulsivity in females ^67–70^, we failed to find an effect on impulsivity in females even when controlling for estrus stage. Patterson et al. looked specifically at the effects of vHPC-ACB inhibition and found subtle sex differences in behavior during motivational conflict ^39^, suggesting that CA1v-ACB inhibition can have differing effects based on sex, which is also consistent with our results.

The vHPC is known to play an important role in regulating food intake control ^23–25,26,27^. While we saw no effect on cumulative food intake or meal patterns with reversible inhibition of the CA1v-ACB pathway, differences between our results and previous findings may be explained by distinct functions of CA1v projection pathways. For example, previous work from our lab reveals that CA1v projections to the lateral hypothalamic area (via ghrelin receptor action) and to the medial prefrontal cortex (via GLP-1 receptor action) potently increases, and reduces food intake, respectively ^52,71–73^. A recent study found that activity of ACB-projecting vHPC neurons increased upon food investigation and that this activity inhibited the transition to meal initiation in mice^74^. However, consistent with our results, chemogenetic manipulation of this pathway in mice did not influence overall food consumption in the long-term ^74^. Thus, our current findings that inhibition of the CA1v-ACB pathway promotes impulsive actions but not food consumption parallels previous work identifying a role for the CA1v-ACB pathway in food anticipatory but not longer-term consummatory behavior, the latter of which is likely driven more by CA1v projections to the lateral hypothalamus and prefrontal cortex.

Considering the significant difference in active lever presses in DRL between control and CA1v->ACB disconnection treatments, we examined whether CA1v-ACB signaling may play a role in willingness to work for palatable food as a readout of food motivation independent from impulsivity. Previous research shows a role for dorsal HPC-ACB circuitry in regulating reward motivation ^75,76^, but the role of vHPC-ACB circuitry is less clear. Yang and colleagues showed that the low frequency photostimulation of vHPC neurons that project to the ACB increases the number of licks per bout – a measure of food palatability – when mice are given access to a sucrose solution, though there was no difference in total bout number – a measure of satiation. Additionally, they revealed that photoinhibition of vHPC neurons that project to the ACB disrupts sucrose-driven flavor preference conditioning ^33^, indicating a role for this pathway in nutritive-based reinforcement learning. This study highlights a role for CA1v-ACB activity in mediating the early ingestive phase positive hedonic experience but reveals little about the effects on effort-based motivation to obtain palatable foods. Thus, our findings that inhibition of CA1v-ACB pathway does not affect operant behavior in the PR task adds to existing literature to suggest that CA1v-ACB circuitry is implicated in behaviors associated with early ingestive phase food reward processes (e.g., food reinforcement-based learning, hedonic evaluations) but does not have a direct effect in an effort-based operant responding for palatable food.

Given that previous research suggests a role for the HPC in guiding anxiety- like behavior^78,79^, we also investigated the effects of CA1v-ACB inhibition on anxiety-like behavior in the zero maze test. Chemogenetic inhibition of the CA1v-ACB pathway had no effect on anxiety-like behavior as measured in this task, consistent with previous pathway-specific vHPC studies ^36,39,80^. Bagot and colleagues found no differences with photostimulation of vHPC-ACB pathway when behavior was assessed using the open field test of anxiety (OFT) ^36^, whereas Patterson et al. also found no differences in anxiety-like behavior when measured using the OFT or the elevated plus maze when inhibiting the vHPC-ACB pathway ^39^. Thus, our results are align with previous reports in revealing little-to-no role of vHPC-ACB circuitry specifically in anxiety-like behavior, collectively indicating that vHPC influences on anxiety likely involve other projection targets.

While prior studies show a role for other pathways originating from the vHPC in shaping impulsive behavior ^22,81,82^, our study suggests, using a novel transsynaptic glutamatergic synapse targeting approach ^46^, that the CA1v-ACB specific pathway plays a role independent of possible collateral projection targets. Additionally, we show that a majority of ACB neurons receiving CA1v projections are inhibitory interneurons. Previous research has shown an important role for inhibitory interneurons in modulating ACB activity, and excitatory dorsal HPC (dHPC) neurons that project to ACB medial spiny neurons (MSNs) also target inhibitory interneurons within the ACB. Thus, present results combined with these previous findings suggest that multiple hippocampal pathways (from both dorsal and ventral subregions) converge on ACB inhibitory interneurons in the ACB to modulate activity in this region. Future studies are required to determine whether vHPC and dHPC inputs to the ACB converge on common vs. distinct neural systems to regulate fundamental behaviors.

In addition to obesity and BED, impulsivity has been implicated in a number of neuropsychiatric disorders, such as compulsive gambling ^9^, substance use disorders ^7,8,83^, and schizophrenia ^84–87^. Similarly, the HPC and ACB have been implicated in many psychiatric conditions, with HPC-ACB circuitry specifically implicated in depression ^35,36,88^, substance use disorders ^89–91^, and schizophrenia ^92–96^. The glutamatergic hypothesis of schizophrenia suggests that hypoactivity of inhibitory interneurons in the corticolimbic circuit may play a crucial role in the development of schizophrenia, particularly parvalbumin+ interneurons ^96–98^, which make up the majority of the interneurons identified in this study. Further, activity of ACB interneurons affects impulsive action ^32^. Future research is needed to investigate the role of ACB-projecting CA1v neurons in schizophrenia pathology, with a focus on whether dysregulation of ACB parvalbumin+ interneurons that receive projections from the Ca1v play a role in disease etiology.

In summary, our results reveal that activity in the CA1v-ACB pathway plays a sex- and task-dependent role in regulating impulsive responding for palatable foods. Calcium-dependent activity in both the CA1v and ACB is significantly elevated prior to a non-impulsive lever press relative to an impulsive press in a test of impulsive action, suggesting a role for both regions in mediating impulse control. Functional testing revealed that chemogenetic inhibition of the Ca1v-ACB pathway – using multiple complementary disconnection approaches – increased impulsive action in males yet had no effect on either impulsive actions or impulsive choices in females. These changes in impulsive action in males are not secondary to changes in food intake, willingness to work for palatable food, or anxiety-like behavior, as these measures were unaffected with pathway inhibition. Future research should further investigate the role of male and female sex hormones in the regulation of this pathway, given the sexual dimorphism observed here. Additionally, characterization of the receptor profiles of both CA1v and ACB neurons in this pathway is warranted, as previous research suggests a role for multiple neurotransmitters, including the dopaminergic ^30,99–101^, opioidergic ^101–104^, and melanin-concentrating hormone ^21^ systems in regulating impulsive responding for palatable foods.

## AUTHOR CONTRIBUTIONS

**Molly E Klug:** Conceptualization, Data curation, Formal analysis, Funding acquisition, Investigation, Methodology, Visualization, Writing – original draft, Writing – review & editing. **Léa Décarie-Spain:** Conceptualization, Investigation. **Logan Tierno Lauer:** Formal analysis, Investigation. **Alicia E. Kao:** Investigation. **Olivia Moody:** Investigation. **Nicholas R. Morano:** Investigation. **Haoyang Huang:** Resources and consultation. **Don Arnold:** Resources and consultation. **Emily E. Noble:** Conceptualization. **Scott E. Kanoski:** Conceptualization, Formal analysis, Funding acquisition, Methodology, Resources, Supervision, Visualization, Writing – review & editing.

---

**Supplementary Figure 1.**
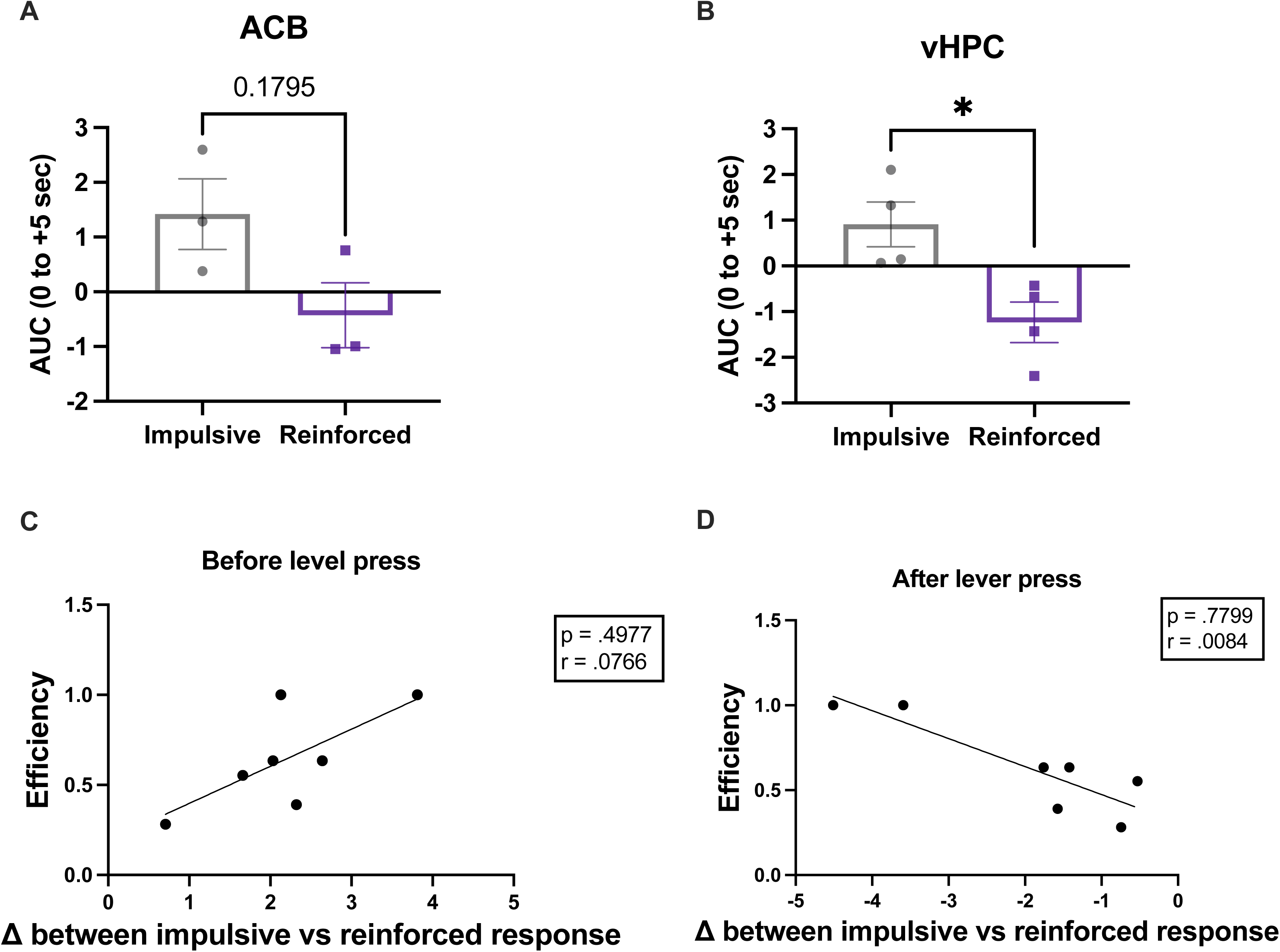
Changes in Calcium-dependent activity is correlated with efficiency in DRL task. Ca^2+^-dependent activity is elevated in the 5s following a non-impulsive lever press, relative to an impulsive press in the ACB **(A)**, and is significantly elevated in the vHPC **(B).** Efficiency in the DRL task is significantly correlated with the change in Ca^2+^-dependent activity both before and after a lever press **(C, D).**

**Supplementary Figure 2.**
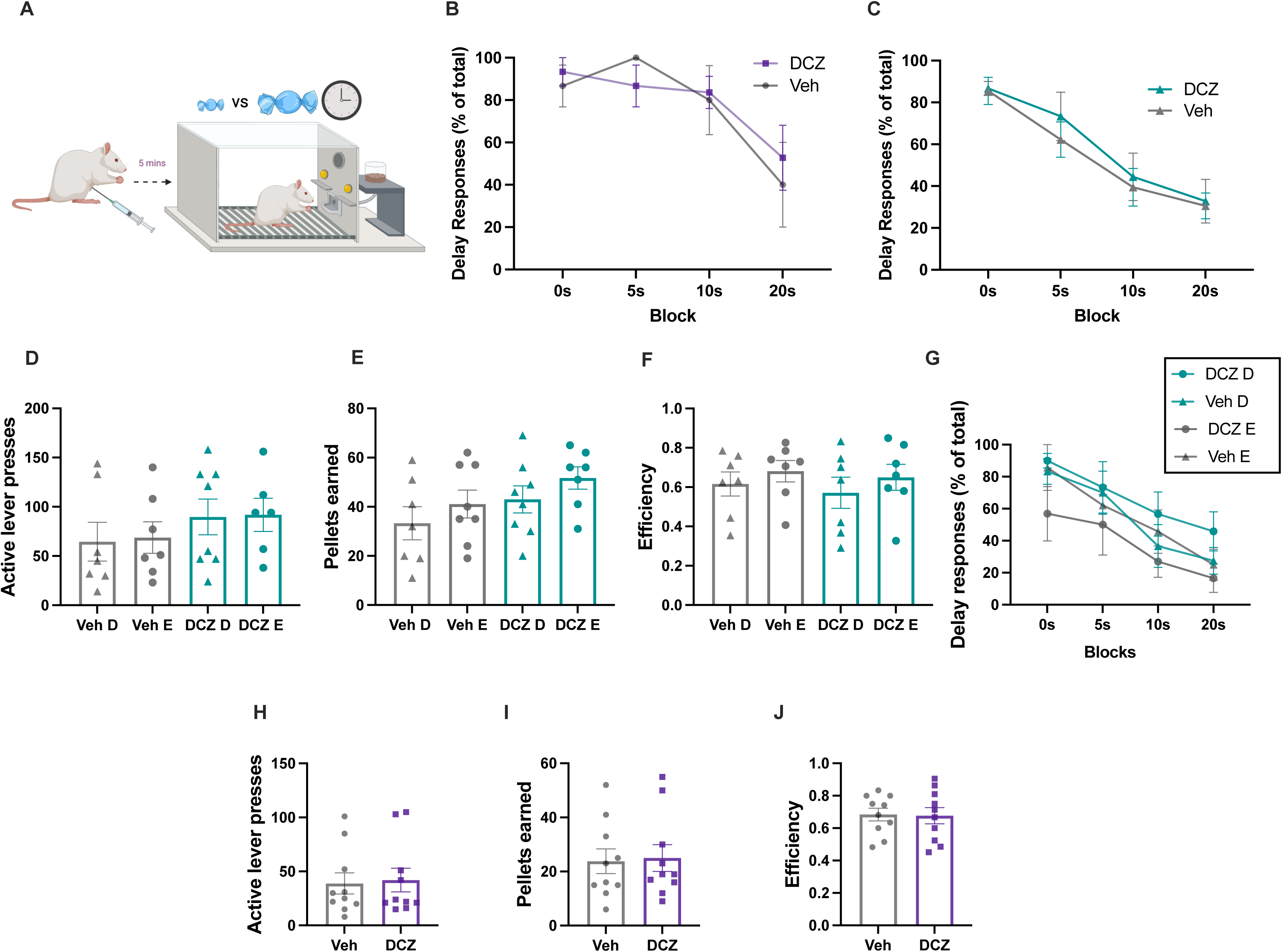
Silencing ACBsh-projecting CA1v neurons does not affect impulsive choice in delay discounting, nor does estrus stage influence impulsive action or choice. **(A)** Diagram depicting the delay discounting task. Chemogenetic inhibition of this pathway does not affect impulsive choice in the delay discounting task in either sex (**B,C)**. Active lever presses **(D),** rewards earned **(E),** and efficiency **(F)** and unaffected by chemogenetic inhibition for female DRL, even when accounting for estrus cycle. Delay discounting is also unaffected by chemogenetic inhibition in females when accounting for estrus cycle **(G)**. Performance in DLR is unaffected by DCZ treatment for DREADDs control group (**H-J**).(DD male n = 6, within-subjects; DD female n = 5-6; between-subjects ; Veh diestrus n = 7, veh estrus n = 7, DCZ diestrus n = 8, DCZ diestrus n = 6 for DRL; between subjects; Veh diestrus n = 6, veh estrus n = 6, DCZ diestrus n = 6, DCZ estrus n = 5 for delay discounting; within subjects; DREADD control n = 10; within-subjects; Data are means ± SEM; *p<0.05).

**Supplementary Figure 3.**
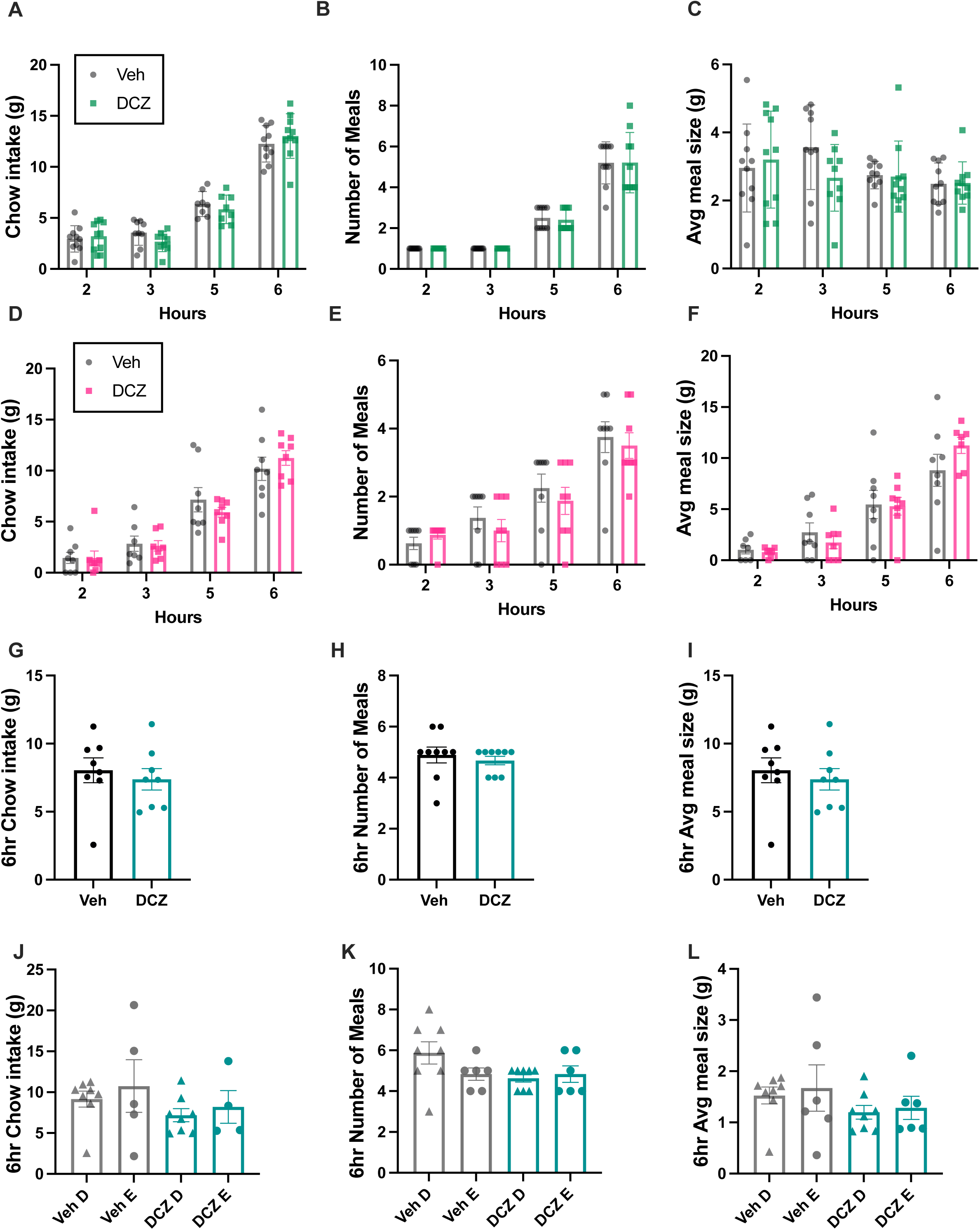
Chemogenetic inhibition of ACBsh-projecting CA1v neurons pathway does not affect home cage intake in males and females. Using both standard chow **(A-C)** and HFHS chow **(D-F)**, chow intake, number of meals, and average meal size is unaffected by DREADDs silencing of vHPC-ACBsh pathway. Cumulative standard chow intake, number of meals, and average meal size is not affected by DREADDs silencing of vHPC-ACBsh pathway in females when analyzed by treatment **(G-I)** or when broken out into estrus stages **(J-L).** (Standard chow males n = 10, HFHS chow n = 8, standard chow females n = 4-8, within subjects for male standard chow and HFHS chow, between-subjects for female standard chow; Data are means ± SEM; *p<0.05, ***p<0.001).

